# The genetic basis of dynamic non-photochemical quenching and photosystem II efficiency in fluctuating light reveals novel molecular targets for maize (*Zea mays*) improvement

**DOI:** 10.1101/2023.11.01.565118

**Authors:** John N. Ferguson, Leonardo Caproni, Julia Walter, Katie Shaw, Min Soe Thein, Svenja Mager, Georgia Taylor, Lee Cackett, Jyotirmaya Mathan, Richard L. Vath, Leo Martin, Bernard Genty, Enrico Pe, Tracy Lawson, Matteo Dell’Acqua, Johannes Kromdijk

## Abstract

Maize (*Zea mays* L.) is a major global crop species which uses C4 photosynthesis. Although C4 is typically considered to be more efficient than C3 photosynthesis, especially under warmer and drier conditions, there is substantial evidence that its efficiency can still be further improved, which may benefit crop performance. Improving photosynthetic efficiency via targeted manipulation of non-photochemical quenching has focused on a limited set of genes that are known to be important determinants of the NPQ response in C3 plants. The C4 pathway may alter NPQ responses but only relatively few studies have explored genetic variation in NPQ kinetics in species that perform C4 photosynthesis. In addition, studies of NPQ responses in field-grown plants of either C3 or C4 species are especially limited. Here we apply high-definition phenotyping of NPQ responses and photosynthetic efficiency and quantitative trait locus (QTL) mapping using a field-grown maize Multi-parent Advanced Generation Inter-Cross (MAGIC) population, which combines the allelic diversity of eight contrasting inbred lines. We find substantial and consistent variation for dynamic NPQ and PSII efficiency for two subsequent field seasons. Further exploration of candidate genes within three major QTL regions identified a strong impact of allelic variation in expression of the minor PSII antenna protein CP24 (LHCB6) on a major QTL for NPQ and efficiency of PSII photochemistry on chromosome 10.

## Introduction

Maize (*Zea mays* L.) is one of the most important cereal crops in the world and is widely cultivated for food, feed, and fuel (Haarhoff and Swanepoel, 2020). Despite this, maize yield trends may be insufficient to meet future demands (Ray et al., 2013). This issue is exacerbated by the increase in frequency and severity of extreme climatic events, such as droughts and heatwaves, due to anthropogenic climate change (Eckardt et al., 2023). Photosynthesis, the process by which solar energy is converted to chemical energy, is the primary source of plant productivity. Consequently, the improvement of photosynthesis has been proposed as an approach through which to increase resilience and yield potential in the crop varieties of the future (Ort et al., 2015).

Maize uses C_4_ photosynthesis, which under warmer and drier environments is more efficient than C_3_ photosynthesis due to a molecular pump that concentrates CO_2_ close to the site of ribulose bisphosphate (RuBP) carboxylase-oxygenase (Rubisco) expression, consequently suppressing RuBP oxygenase activity and associated energetic losses via the photorespiration pathway (Hatch, 1987). Despite these benefits of the C_4_ photosynthetic pathway, improving photosynthetic efficiency in maize may also lead to increased productivity (Salesse-Smith et al., 2018) and potentially mitigate environmental stress (Doron et al., 2020; Salesse-Smith et al., 2020). Increasing photosynthetic efficiency could therefore be an important trait to maintain current and future maize productivity (see also review by Sales et al., (2021))

Excessive light energy absorbed by the photosynthetic antennae enhances the probability of formation of reactive oxygen species (ROS). ROS can inflict oxidative damage to thylakoid components such as the PSII reaction center protein D1 and thereby lead to deactivation of photochemistry (Aro et al., 1993). This so-called photoinhibition gives rise to sustained downregulation of photosynthetic efficiency and can therefore reduce crop carbon gain and associated growth and productivity (Long et al., 1994). To reduce the risk of photoinhibition, photoprotective non-photochemical quenching (NPQ) routes are induced under high light, which dissipate excess energy in a controlled manner (Murchie and Ruban, 2020). The sporadic nature of light availability in crop canopies (Long et al., 2022) means that rapid induction and relaxation responses of NPQ are necessary to adequately adjust the efficiency of light-harvesting to the intensity of intercepted light. Model simulations (Zhu et al., 2013) and proof of concept studies in field-grown tobacco and soybean (Kromdijk et al., 2016; De Souza et al., 2022) suggest that accelerating NPQ relaxation can enhance crop yields by reducing the lag-time to return to high PSII operating efficiency and CO_2_ fixation upon a switch from high to low light. In addition, enhancing the NPQ amplitude and rate of induction have also been linked with increased yields in rice (Hubbart et al., 2018). These proof-of-concept studies have focused on a limited set of genes that are known to be important determinants of the NPQ response in C_3_ plants, namely PSII subunit S (PSBS), violaxanthin de-epoxidase (VDE) and zeaxanthin epoxidase (ZEP). Only relatively few studies have explored NPQ kinetics in species that perform photosynthesis via the C_4_ pathway, which may alter NPQ responses (Guidi et al., 2019). In addition, studies of NPQ responses in field-grown plants of either C_3_ or C_4_ species are especially sparce.

Mapping populations and intraspecific diversity panels offer powerful tools to accelerate the identification of genetic determinants underpinning natural diversity for photosynthetic efficiency and induction and relaxation responses of NPQ, yielding allelic variants that may have immediate breeding relevance. To date, the authors are aware of only one recent study (Sahay et al., 2023) that evaluated the genetic determinants of variation in NPQ responses under field conditions (US Midwest, Nebraska) across a diverse panel of maize accessions. Ample variation was found, yielding several putative quantitative trait loci (QTL) based on genome-wide association analysis (GWAS). Here, we aimed to expand on the results by Sahay et al. by developing an enhanced understanding of the genetics underpinning dynamic photoprotection and PSII operating efficiency in maize in a maritime climate close to the northern latitude edge of maize cultivation, where maize acreage has expanded substantially in recent decades (UK government, 2023). To do so, we characterized NPQ responses and photosynthetic efficiency across a Multiple parent Advanced Generation Inter-Cross (MAGIC) maize panel (Dell’Acqua et al., 2015), which combines the allelic diversity of eight maize inbred lines enabling high power and high-resolution mapping of complex traits (Scott et al., 2020). The results showed substantial and consistent variation for dynamic NPQ and PSII efficiency leading to identification of QTL. Using a combination of genomics, transcriptomics, protein biochemistry and targeted physiological phenotyping, putative causal genes were explored within three major QTL regions.

## Results

### Genetic variation for PSII efficiency and NPQ across the maize MAGIC population

320 MAGIC maize recombinant inbred lines (RILs) were grown in a replicated alpha-lattice design in the east of the United Kingdom (52.2 °N, 0.1 °E) for two consecutive field seasons (2021 and 2022). We phenotyped 315 RILs in 2021 and 312 in 2022, where 301 RILs were common between the two years (Supplemental Table S1-2). This represented data collected from more than 3700 leaves in total.

A chlorophyll fluorescence protocol was employed to determine timeseries data for photosystem II efficiency (ΦPSII) and NPQ induction and relaxation in response to 10 min high light exposure followed by 12 min dark recovery. To measure all genotypes in high throughput to sample all genotypes, the assay was performed on leaf segments, which we have recently demonstrated to give the same results as intact leaves still attached to the plant (Ferguson et al., 2023a). The resulting NPQ and ΦPSII values were analyzed separately for each measured timepoint as well as modelled across the light and dark phases of the protocol (Figure 1).

**Figure 1.**
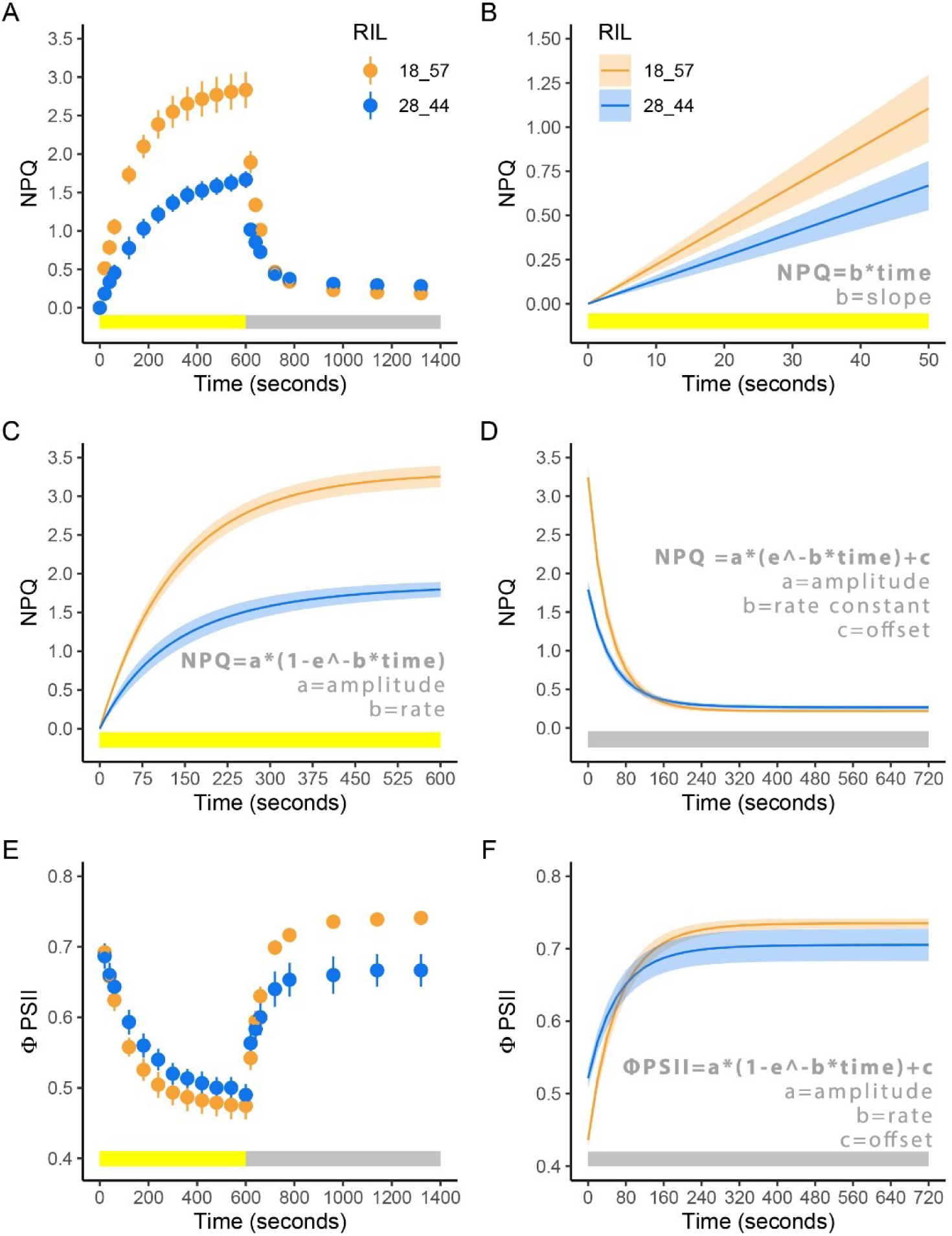
Example of models used to fit the dynamic NPQ and ΦPSII data based on two distinct recombinant inbred lines (RILs; SSA_00157 and SSA_00376). Data shown are from six biological reps of each RIL. (A) NPQ throughout the measurement protocol. Circles represent the mean value, and the bars represent the standard error of the mean. (B) Initial NPQ induction modelled via a linear regression. The solid line represents the mean of the predicted NPQ according to the linear regression fit and the ribbon represents the standard error of the predicted fit. (C) NPQ induction modelled via an exponential equation. (D) NPQ relaxation modelled via an exponential model. (E) ΦPSII through the measurement protocol. (F) ΦPSII recovery modelled via an exponential model.

Substantial genetic variation was detected for all measured and modelled traits in both years (Figure 2; Supplemental Figure S1; Supplemental Tables S1-2). The 2022 growing season was drier and warmer than the 2021 growing season and necessitated irrigation (Supplemental Figure S2). Consistent with the drier and warmer conditions in 2022, the population average for maximum NPQ was higher and the rates of relaxation of NPQ and recovery of ΦPSII following the actinic light being switched off were slower than in 2021 (Figure 2). Despite these environmental differences, the population-wide range of variation was similar across both experimental years and strong year by year correlations were found (Figure 2; Supplemental Figure S1), indicative of a strong genetic basis to the traits in question. Indeed, broad-sense heritability across all traits (H^2^_B_) computed both for each year individually and jointly, ranged between 0.3-0.7 (Supplemental Figure S3).

**Figure 2.**
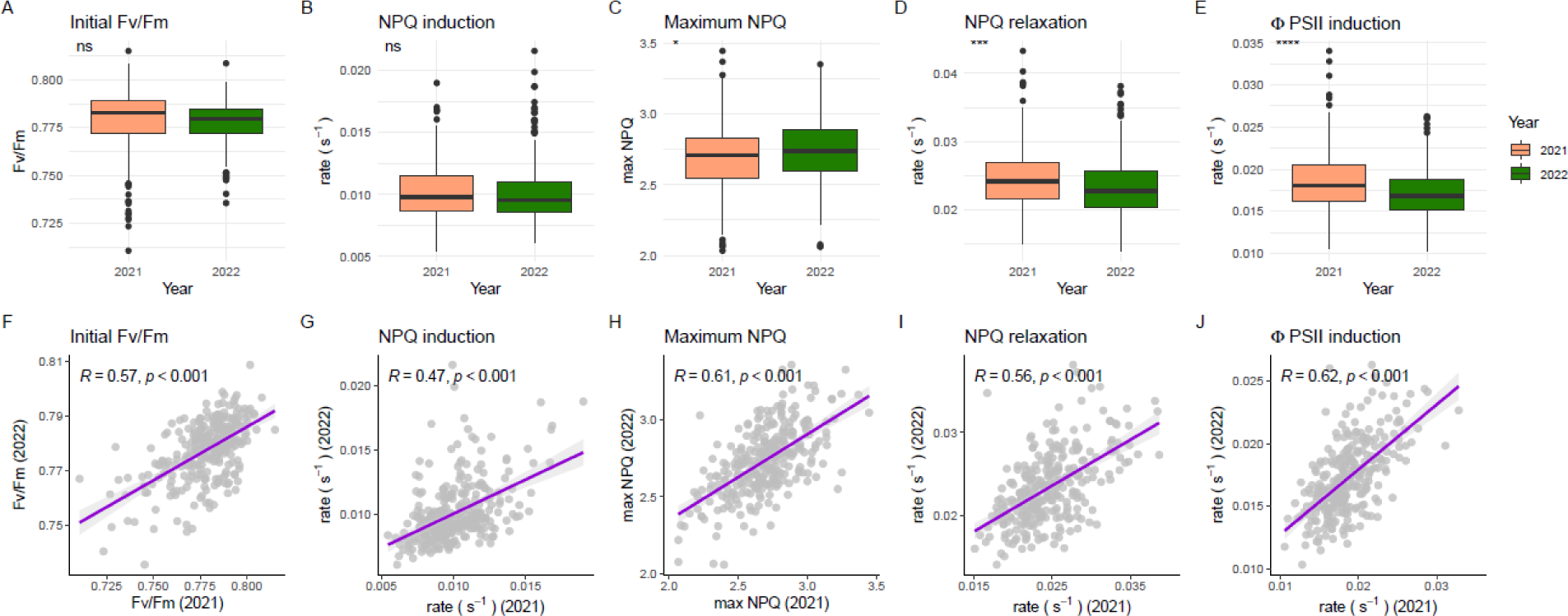
Comparison of selected parameters across the two experimental years. (A-E) Boxplots demonstrating population wide variation across both experimental years for Fv/Fm, NPQ induction rate, maximum NPQ, NPQ relaxation rate, and ΦPSII recovery rate. (F-H) Scatter plots demonstrating associations between the same traits across each experimental year. Significant differences and correlations are denoted at the following *p*-value levels: * 0.05, ** 0.01, *** 0.001, **** 0.0001.

Pair-wise trait correlations were largely conserved across the 2021, 2022, and jointly predicted accession means. If a pair-wise trait correlation was significant in 2021 it was also significant in 2022 and vice versa for 75 of the 91 interactions tested (Supplemental Table S3). Interestingly, significant negative associations between the rates of NPQ induction and relaxation were observed across both years, suggesting that RILs with fast rates of induction of NPQ had slower rates of relaxation and vice versa. Since the NPQ relaxation rate positively correlated with the rate of recovery of ΦPSII, the latter was also observed to have a significant negative correlation with the rate of NPQ induction across both years (Supplemental Table S3).

### Multiple QTL define variation in PSII efficiency and NPQ and demonstrate differential importance depending on the light environment

74,706 high-quality single nucleotide polymorphism (SNP) markers obtained from Single Primer Enrichment Technology (SPET; (Scaglione et al., 2019)) on the founder lines and RILs were used to probabilistically reconstruct the RIL genomes *in silico* and to perform QTL mapping on the basis of parental haplotypes. We first focused on identification of QTL for NPQ and ΦPSII at each timepoint throughout the measurement protocol, based on predicted means derived from the joint year model. The QTL mapping results for these traits showed that the genomic regions associated with the observed variation were dependent on the time elapsed following the actinic light being switched on or off (Supplemental Movie S1-4). For example, variation in NPQ during the light induction phase of the experimental protocol was initially defined by a QTL on chromosome 10 (Figure 3A) which rapidly disappeared before QTL on chromosomes 9, 5, and 1 sequentially appeared (Figure 3B-C). Similarly, variation in NPQ during the dark relaxation phase was initially associated with the same chromosome 1 QTL as the light phase (Figure 3D), before other QTL on chromosomes 2 (Figure 3E) and 6 (Figure 3F) passed the significance threshold. The genetic architecture of ΦPSII during the assay was relatively less complex. A highly significant large-effect QTL for dark-adapted ΦPSII (Fv/Fm) was identified on chromosome 10 co-located with the induction phase NPQ QTL (shown in Figure 3A). This QTL persisted for most time-points during the light phase (Supplemental Figure S4A), where ΦPSII rapidly declined as well as during the dark recovery phase, gradually increasing in significance (Supplemental Figure S4C-D, Supplemental Movie S3-4). An additional QTL for ΦPSII during the light phase was found on chromosome 9 for time-points from 60 s onwards (Supplemental Figure S4B).

**Figure 3.**
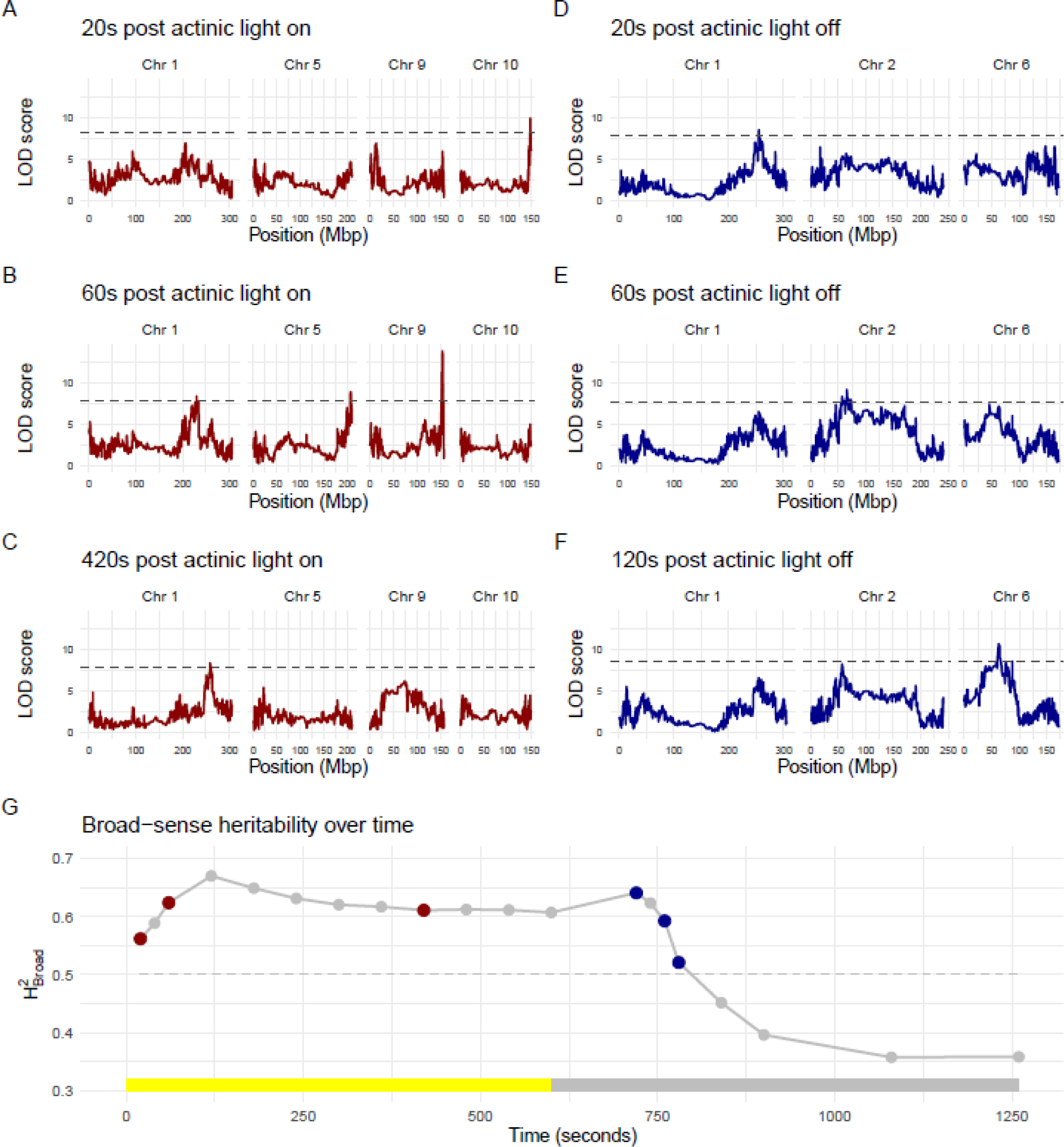
QTL mapping of NPQ at different timepoints within the induction (light) and relaxation (phases). Mapping is performed using the predicted means derived from the joint-year linear mixed effect models. (A) QTL for NPQ 20 seconds after the actinic light is switched on. (B) QTL for NPQ 60 seconds after the actinic light is switched on. (C) QTL for NPQ 420 seconds after the actinic light is switched on. (D) QTL for NPQ 20 seconds after the actinic light is switched off. (E) QTL for NPQ 60 seconds after the actinic light is switched off. (F) QTL for NPQ 120 seconds after the actinic light is switched off. (G) Broad-sense heritability (H^2^_B_) for NPQ over time.

The same QTL were also found when mapping the fluorescence parameters that were determined from combined analysis of several time-points, reducing the risk of spurious associations based on only one parameter or timepoint. For example, the QTL on chromosome 10 for both NPQ and ΦPSII was also detected for PI and for the y-axis intercept from the exponential model describing the recovery of ΦPSII. This reflects the close correlations between these traits and their common association with PSII photochemistry (Supplemental Table S3). The rate constant of NPQ induction was estimated by the initial slope of NPQ during the first 60 seconds of the light phase as well as by a first-order exponential model for the whole light phase, both of which were associated with the QTL on chromosome 9 previously found for several early timepoints during the light phase (Table 2). Interestingly, the QTL on chromosome 1 found later in the light phase and early dark phase, was also found for the amplitude of NPQ induction, amplitude of NPQ relaxation and amplitude of ΦPSII recovery. This is consistent with the mechanistic links between NPQ relaxation and recovery of ΦPSII following a change in light intensity (Table 2).

### Candidate gene prioritization

Altogether, all QTL identified contained 3064 unique gene models (Supplemental Table S4). For all QTL-associated gene models we identified the most similar Arabidopsis ortholog and used gene ontology to find genes with known roles in photosynthesis and in the detoxification of reactive oxygen species (summarized in Supplemental Figure S5). However, exclusive prioritization based on gene ontology annotations would rule out discovery of genes with no prior association to the phenotypes measured here. We therefore employed several additional analyses to further prioritize candidate genes underpinning the QTL on chromosome 1, 5, 9 and 10.

#### Expression level differences in VTE4 correlate with QTL effects on NPQ on chromosome 5

First, assuming that a difference in gene expression may drive observed phenotypic differences, expression levels of QTL-associated gene models in the founder lines were analyzed for significant correlations with the estimated contribution of founder haplotypes to the QTL. Altogether, through this approach, 46 unique genes were identified which had constitutive differences in expression levels that were correlating with clustered founder effects on QTL (Supplemental Table S5). Clustering the founder effects in the QTL confidence interval of the chromosome 5 QTL for NPQ induction (Figure 3B, 5A) showed a negative effect on the QTL (lower NPQ) associated with the founder haplotype from HP301, while all the other founder haplotypes were clustered around neutral effects (Figure 4B). We found that of the 341 protein-coding genes in the QTL confidence interval (Supplemental Table S4), expression levels of six genes significantly correlated with these founder effects. These six genes included the gamma-tocopherol methyltransferase gene *VTE4* (Figure 4C), which is involved in the biosynthesis of tocopherols, a major class of antioxidants with well-known roles in photoprotection (Havaux et al., 2005).

**Figure 4.**
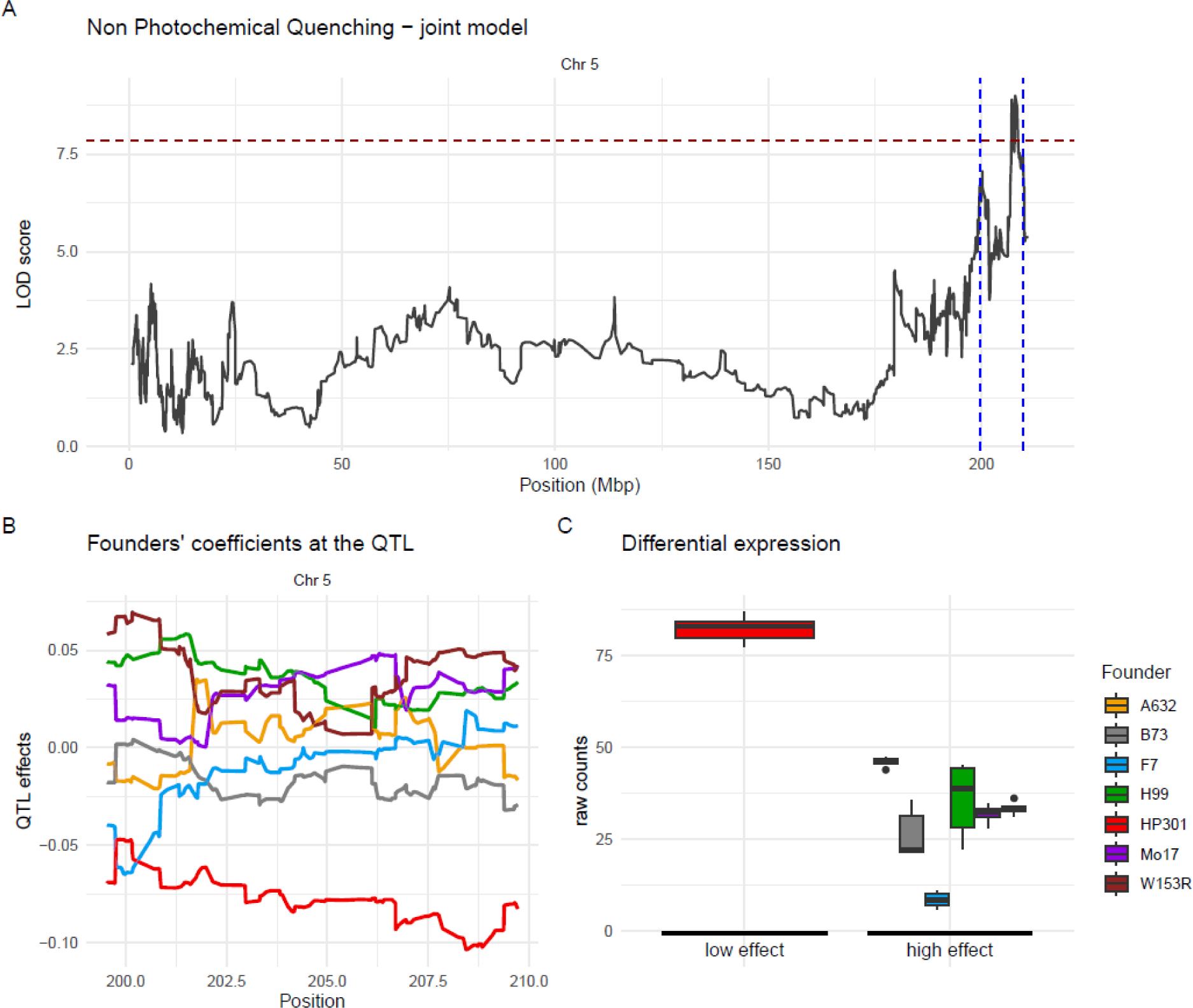
Association of VTE4 expression variation with founder effects at QTL for NPQ. (A) QTL results for NPQ 60s post actinic light being switched on. The confidence intervals of the QTL are denoted by the blue dashed lines. (B) Effects of each founder at the QTL. (C) Expression difference of VTE4 across the founders grouped according to the QTL effect clustering.

#### A large-effect QTL on chromosome 9 is a key regulator of NPQ and Φ_PSII_ during early induction

A large-effect QTL on chromosome 9 was detected for both NPQ and Φ_PSII_ based on both linear and exponential models (Figure 5A; Table 1) as well as individual timepoints up to 120 s (Table 2). The B73 founder haplotype seemed to have strong opposite effects for this QTL: positive for NPQ and negative for ΦPSII. Two genes within the confidence intervals of this QTL had expression patterns matching the QTL effect estimates (Supplemental Table S5): the closest Arabidopsis orthologs of these genes are a leucine-rich repeat protein kinase and a CCT motif protein, both remain currently uncharacterized.

**Figure 5.**
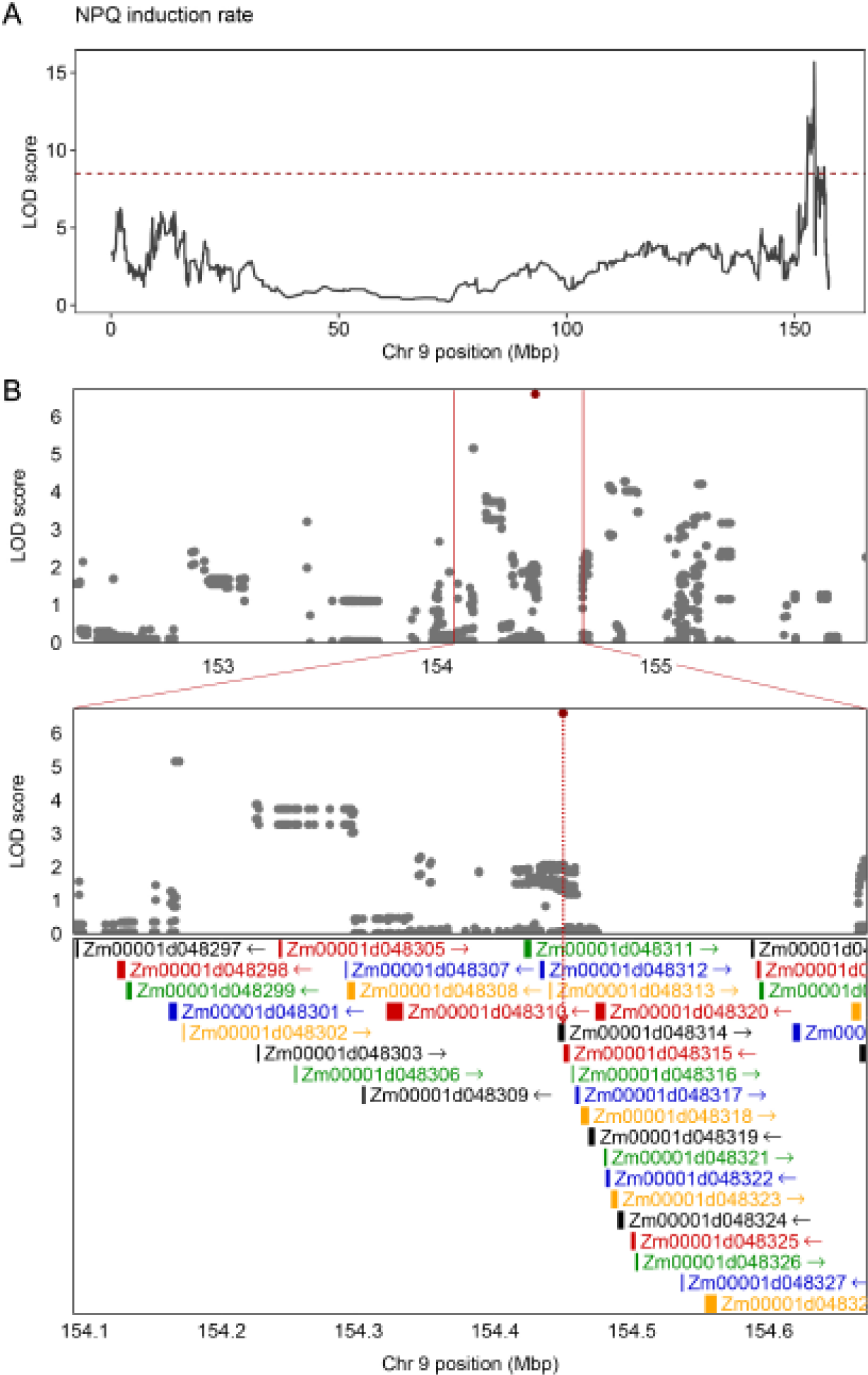
(a) LOD score plot of NPQ induction rate on Chromosome 9 QTL, the horizontal dotted line represents the threshold of significance. (b) top panel: SNP association mapping within a 4 Mbp region, centered on NPQ induction rate QTL on chromosome 9. Bottom panel: Zoomed in view of the same region (± 0.25 Mbp, centered on QTL), showing a putatively associated SNP falling within the coding region of Zm0001d048314. In this panel all 36 gene models annotated in this zoomed-in region are shown. In both panels, SNPs having LOD scores > 99^th^ percentile are highlighted in red.

**Table 1.** Quantitative trait loci (QTL) associated with NPQ and ΦPSII at each measured timepoint.

| Period | Model | Phenotype | Chr. | Average Peak Position<br>(Mbp) | max LOD | min CI- low<br>(Mbp) | max CI- high<br>(Mbp) | Time (seconds) |
| --- | --- | --- | --- | --- | --- | --- | --- | --- |
| Dark | Joint-year | NPQ | 1 | 255 | 8.5 | 247 | 262.1 | 20 |
| Dark | Joint-year | phiPSII | 10 | 149.2 | 19.9 | 147.8 | 150 | 20, 40, 60, 120, 180, 360, 540 |
| Light | Joint-year | NPQ | 10 | 148.7 | 9.9 | 147.7 | 149.5 | 20 |
| Light | Joint-year | NPQ | 9 | 154.4 | 15.6 | 152.9 | 154.5 | 40, 60, 120 |
| Light | Joint-year | NPQ | 5 | 208.1 | 9 | 199.5 | 209.7 | 60, 120 |
| Light | Joint-year | NPQ | 1 | 258.4 | 8.5 | 248.3 | 263.5 | 240, 300, 360, 420, 480, 540, 600 |
| Light | Joint-year | NPQ | 1 | 229.8 | 8.5 | 204.5 | 234.3 | 60 |
| Light | Joint-year | phiPSII | 10 | 149.1 | 21.3 | 147.8 | 149.5 | 20, 40, 60, 180, 240, 300, 360, 420, 480, 540, 600 |
| Light | Joint-year | phiPSII | 9 | 154.4 | 11.6 | 154 | 155.4 | 60, 120 |
| Dark | 2021 | phiPSII | 10 | 149.18 | 18.97190182 | 147.739499 | 149.953864 | 20, 40, 60, 120, 180, 360, 540 |
| Light | 2021 | NPQ | 10 | 148.67 | 11.3 | 146.3 | 149 | 480 |
| Light | 2021 | NPQ | 9 | 154.36 | 13.7 | 152 | 154.7 | 40, 60, 120 |
| Light | 2021 | NPQ | 5 | 207.76 | 10.9 | 195.5 | 210 | 60, 120 |
| Light | 2021 | NPQ | 3 | 158.56 | 8.5 | 150.3 | 164.5 | 240, 300, 360, 420, 480, 540, 600 |
| Light | 2021 | phiPSII | 10 | 149.03 | 20.6 | 148.5 | 149.5 | 20, 40, 60, 240, 300, 360, 420, 480, 540, 600 |
| Light | 2021 | phiPSII | 9 | 154.36 | 10.7 | 154.0 | 155.6 | 60, 120 |
| Dark | 2022 | NPQ | 2 | 240.57 | 10.4 | 240.3 | 241.9 | 180 |
| Dark | 2022 | NPQ | 1 | 249.32 | 10.8 | 215.1 | 260.9 | 120, 180 |
| Dark | 2022 | NPQ | 1 | 229.83 | 8.6 | 215.1 | 234.0 | 60 |
| Dark | 2022 | phiPSII | 10 | 149.18 | 12.6 | 148.0 | 149.5 | 120, 180, 360, 540 |
| Light | 2022 | NPQ | 9 | 154.40 | 11.8 | 136.3 | 155.2 | 60, 120, 180 |
| Light | 2022 | phiPSII | 10 | 149.18 | 12.6 | 148.0 | 150.0 | 20, 50, 480, 540, 600 |
| Light | 2022 | phiPSII | 9 | 154.49 | 10.6 | 154.4 | 155.3 | 120, 180 |

In addition to these two gene models, a few other candidate genes stood out based on their functional annotation. In particular, the QTL confidence interval was found to contain the *Carotenoid cleavage dioxygenase 1 (CCD1)* gene. In maize, extensive *CCD1* tandem copy number variation has been documented to underpin the *White Cap* (*Wc*) effect, which is prevalent in white-grain varieties where it leads to white endosperm via enhanced carotenoid break-down (Tan et al., 2017). Based on the well-known role of carotenoids in ROS scavenging (Domonkos et al., 2013), we tested whether *CCD1* copy number variation was present across the founders of the MAGIC population that might correspond to their effects at this QTL. However, since all major founders were found to have just one copy of CCD1, (with the exception of CML91 which had 10 copies but only contributes <5% of total allelic variation), this hypothesis could be discarded (Supplemental Table S6). The absence of any structural variation like presence of CNV could also be confirmed using normalized Whole Genome Sequencing (WGS) read counts of the founders at QTL region (Supplemental Table S7).

Finally, we projected whole-genome imputed SNPs of the MAGIC founders onto the RILs leveraging the haplotype reconstruction of their genomes at this locus. 5324 SNP markers contained within the 4 Mb QTL interval were used for association mapping using a standard mixed linear model with kinship correction (Figure 5B). The most significant SNP arising from this approach was found within the *Zm0001d048314* gene model, which according to PLAZA Integrative Orthology (Van Bel et al., 2018) encodes a chloroplastic protein kinase matching an orthologous gene family of Arabidopsis consisting of 45 predominantly PBS1-Like kinases (Supplemental Table S8). Amongst these is the PROTEIN KINASE 1B gene (PK1B or APK1B), which has been implicated in light-induced stomatal opening (Elhaddad et al., 2014) and could indirectly be related to our identification of this QTL for NPQ and ΦPSII during the light-phase of the protocol.

#### A deficient CP24 allele may explain the large effect QTL for NPQ and ΦPSII on chromosome 10

Finally, we focused on the large effect QTL in the distal portion of chromosome 10 which was found across many of the ΦPSII and NPQ parameters (Figure 3; Supplemental Figure S4, Table 1&2). For illustrative purposes, here we show data from the joint-year model for Fv/Fm (Figure 6A). The QTL effects estimated per founder haplotype clearly showed that this QTL was driven by a negative effect associated with the F7 haplotype (Figure 6B). From the 188 gene models within the QTL confidence interval (Supplemental Table S4), expression levels of 10 genes were significantly correlated with the founder QTL effects (Supplemental Table S5). These genes included the minor PSII antenna *CP24* (lhcb6) expression of which was significantly reduced in F7 based on both RNA-seq data representing all founder lines, as well as based on independent expression data for F7 and B73 (Figure 6C) from Cackett et al., (2023) Since *CP24* is known to affect the structure and function of the PSII light harvesting antenna in Arabidopsis (Kovács et al., 2006), we decided to further explore a causative link with the QTL effects observed here. Immunoblot analysis of B73 and F7 using a CP24-specific antibody confirmed that F7 had severely reduced CP24 protein abundance compared to B73 (Figure 6F), which was consistent with a decreased abundance of higher order PSII-LHCII super complexes in blue native PAGE analysis of purified thylakoid membranes from both founder lines (Figure 6G) indicating significant truncation of PSII antenna size associated with the F7 haplotype. We confirmed the reduction in *F*_v_/*F*_m_ in F7 relative to B73 (Figure 6D). Moreover, the function of PSII antenna size was further assessed *in vivo* using fast fluorescence kinetics (Supplemental Figure S6), showing significantly lower fluorescence rise times in F7 relative to B73 (Figure 6E) with a lower Fv/Fm mainly associated with an increase in Fo (Supplemental Figure S6), which is indicative of an alteration of light harvesting efficiency and photochemical yield (de Bianchi et al., 2008; Kovács et al., 2006; Ilíková et al., 2021). Altogether, these results are consistent with allelic variation in *CP24* to contribute to the QTL effect on chromosome 10.

**Table 2.**
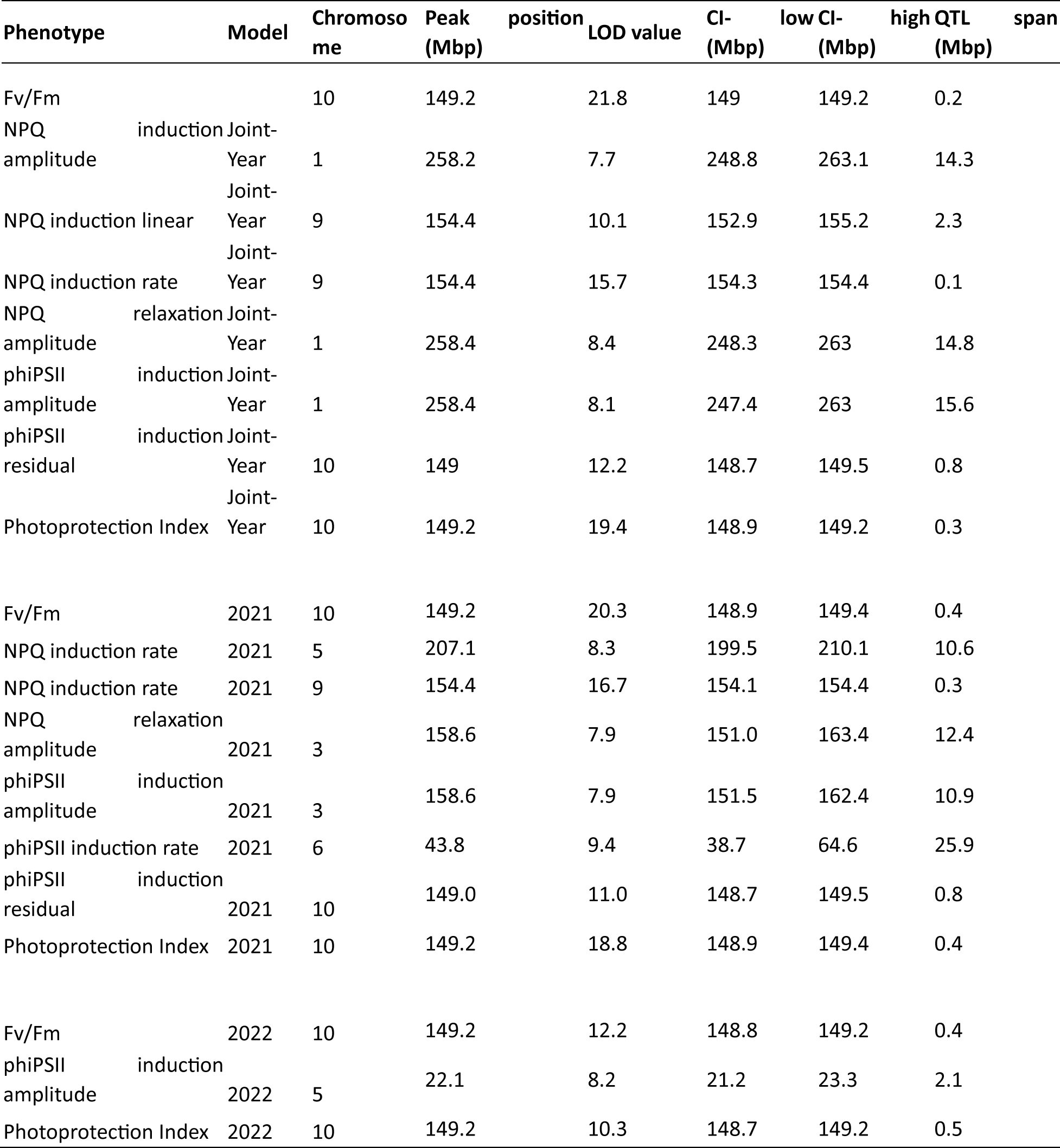
Quantitative trait loci (QTL) associated with Fv/Fm and traits derived from the linear and exponential modelling of NPQ induction, NPQ relaxation and ΦPSII recovery.

**Figure 6.**
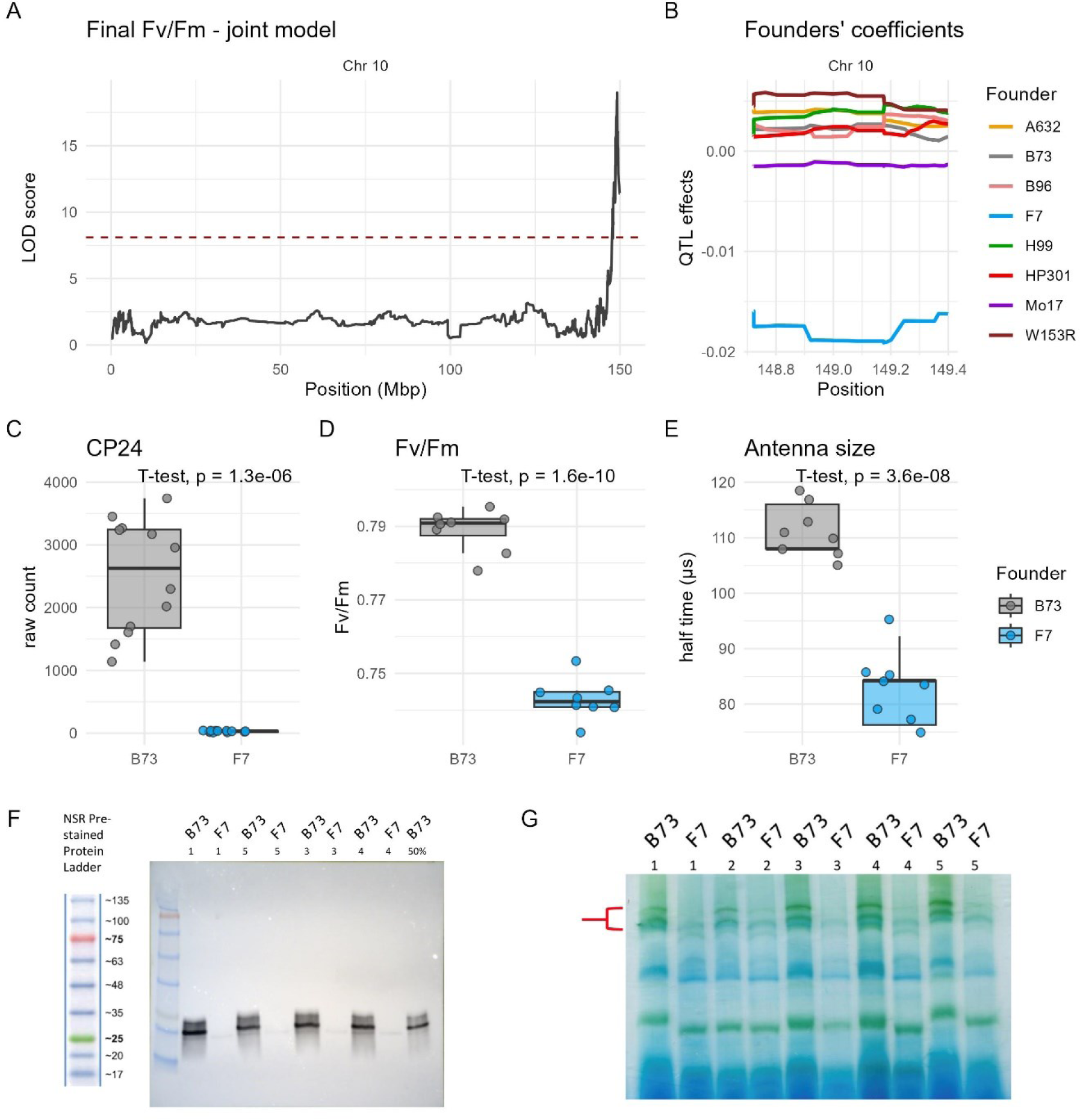
(A) Chromosome 10 QTL for Fv/Fm (B) Founder QTL effects at chromosome ten QTL (C) Difference in expression of CP24 between B73 and F7 from Cackett *et al*. (2023). (D) Difference in *F*_v_/*F*_m_ assessed before the OJIP induction curve. (E) Estimation of function antenna size based on half time to maximal fluorescence during OJIP fluorescence induction curve. (F) Western blotting of B73 and F7 thylakoid membrane protein samples using a CP24-specific antibody. (G) Blue native gel electrophoresis (BN-PAGE) of thylakoid membrane complexes of B73 and F7.

## Discussion

Natural genetic variation in photosynthetic traits provides the starting point for non-transgenic strategies aiming to achieve improved photosynthetic efficiency in crops (Lawson et al., 2012). Here we used a high-throughput protocol to characterize induction and relaxation of NPQ and ΦPSII across 311 maize MAGIC RILs and 9 founder lines grown close to the northern latitude edge of maize cultivation (52.2 °N, 0.1 °E). The results confirm the existence of substantial heritable variation across the measured or calculated traits. Below we discuss the implications of our findings for the genetic architecture of these traits in MAGIC populations, strategies to prioritize candidate genes and utility for improving photosynthetic efficiency in maize.

### The genetic architecture of ΦPSII and NPQ traits reflects high power and tractable complexity of marker-trait associations in the MAGIC maize population

The kinetics of NPQ induction and relaxation can have a strong impact on the performance of photosynthesis under fluctuating light conditions, which is increasingly recognized as an important determinant of photosynthetic efficiency under field conditions (Long et al., 2022; Kaiser et al., 2018) and natural genetic variation has been suggested to hold promise for improvement strategies (Murchie et al., 2018). Despite this, only a few studies have evaluated phenotypic variation in induction and relaxation kinetics of NPQ and ΦPSII across large field-grown populations to identify genetic associations. (Wang et al., 2017) measured NPQ levels on a diverse collection of field-grown rice accessions leading to GWAS-based identification of 33 unique associated SNPs, some of which were validated using bespoke F2 populations. (Sahay et al., 2023) characterized the genetic determinants of ΦPSII and NPQ traits in a maize diversity panel with 751 accessions grown in the US Midwest (Lincoln, NE), yielding 18 unique SNPs significantly associated with at least one trait and an additional 185 unique SNPs associated at a lower significance threshold, demonstrating a multitude of small-effect genetic loci. Trait broad-sense heritability and range of variation were similar between (Sahay et al., 2023) and the current study. However, here QTL mapping of ΦPSII and NPQ by time-point, as well as of parameters derived from multiple timepoints consistently identified two large effect QTL shared between the joint year models for ΦPSII and NPQ, and three additional QTL specifically for NPQ. This reduced complexity in genetic architecture compared to Sahay *et al*. (2023) reflects the contrasting population used here, where the strength of the MAGIC population design resides in the high power of QTL detection based on intermediate allelic diversity across the founders. Although mapping resolution in diversity panels such as used by Sahay et al. (2023) may benefit from fast linkage disequilibrium (LD) decay (Yan et al., 2009), a MAGIC population has the advantage of low population structure and fully resolved identity by descent (Dell’Acqua et al., 2015). The identification of causal genes underlying QTL identified through either linkage mapping or GWAS remains a significant challenge for genetics (Bergelson and Roux, 2010). Even in situations in which genomes are well annotated and gene ontology information is available, it can still be difficult to pinpoint causal genes with some confidence, as the extent of linkage disequilibrium combined with less-than-complete molecular marker coverage of the target genomes limits the definition of the mapping exercise (Lin et al., 2019). The definition of QTL mapping is increased in multiparent populations (Scott et al., 2020) such as used here, yet most QTL still contain tens or hundreds of genes (Table 1-2), necessitating prioritization in the selection of causal variants for further evaluation. Identification of candidate genes across multiple environments or populations increases confidence in a potential causal role. It was therefore interesting to find a total of 20 unique genes within QTL confidence intervals from both this study and (Sahay et al., 2023) Supplemental Table S9 and S10), none of which has previously demonstrated roles in photosynthesis.

### Prioritization of candidate genes based on gene expression data

One potential means for genetic variants to give rise to QTL effects is via impacting transcription of genes via variation in proximal or distal regulatory elements. Transcriptome-wide association study (Gamazon et al., 2015) has been developed to identify transcriptionally regulated genes in association with complex traits, which in combination with GWAS or QTL mapping can help prioritize causal genes by means of colocalization, as demonstrated recently for water use efficiency trait variation in *Sorghum bicolor* (Ferguson et al., 2021). Here, we used pre-existing expression data for the founder lines specifically for the gene models contained within each QTL interval to test for significant correlations between gene expression level variation and QTL effects by founder haplotype. This strategy filters for genes which are consistent with the case where a genetic variant associates with both the expression level of a causal gene and the trait of interest. Although it should be emphasized that this is only one non-exclusive mode of action to explain QTL effects, results from this strategy included three genes where prior knowledge of their functional role seemed consistent with the observed QTL effects.

Firstly, the vitamin E biosynthesis gene *VTE4* sequence resides within the QTL for NPQ induction on chromosome 5 and founder variation in expression levels of *VTE4* showed a significant negative correlation with the clustered founder QTL effects. The gene product of *VTE4* is a gamma-tocopherol methyltransferase (γ-TMT), which controls the synthesis of α-tocopherol from γ-tocopherol. Tocopherols are lipophilic antioxidants and α-tocopherol accumulates predominantly in photosynthetic tissue (DellaPenna and Pogson, 2006), where it seems to act as a ROS scavenger to protect thylakoid pigments, proteins as well as membrane lipids. When grown outside, *vte-4* mutant lines in *Arabidopsis thaliana* which are deficient in α-tocopherol and constitutively accumulate γ-tocopherol, showed significant reductions in chlorophyll and carotenoid content (Semchuk et al., 2009). In addition, overexpression of γ-TMT may be beneficial for abiotic stress tolerance (Ma et al., 2020). Interestingly, there is some evidence that α-tocopherol and zeaxanthin have partially overlapping antioxidant roles (Havaux et al., 2005). The *npq1* mutant in *A. thaliana*, which lacks violaxanthin de-epoxidase and therefore the ability to form zeaxanthin, accumulates significantly more α-tocopherol than wildtype plants (Havaux et al., 2000) and vice versa, tocopherol-deficient *vte1* plants grown under high light contained more zeaxanthin (Havaux et al., 2005). Zeaxanthin functions both as a ROS scavenging molecule and as a positive regulator of NPQ. Therefore, if the different *VTE4* expression levels in the MAGIC RILs lead to similar functional redundancy patterns between zeaxanthin and α-tocopherol observed in *A. thaliana* mutants, these may explain the NPQ effect of this locus.

Secondly, the shared large-effect QTL for ΦPSII and NPQ on chromosome 10 contains a maize CP24 ortholog for which expression levels were significantly positively correlated with the QTL effects on both traits. CP24 (*LHCB6*) is a monomeric light-harvesting complex protein which is positioned between the moderately bound LHCII trimers and the core complex. As with the other minor antennae CP26 and CP29, CP24 is essential for efficient energy transfer from the distal LHCII to the core complex (Arshad et al., 2022). CP24 seems to have evolved in land plants and has been lost in some subgroups of gymnosperms (Koziol et al., 2007; Kouřil et al., 2016). *A. thaliana* knock-out mutants of CP24 show a pronounced reduction of maximal ΦPSII, NPQ amplitude and antenna supercomplex abundance (de Bianchi et al., 2008; Kovács et al., 2006). The chromosome 10 QTL effect detected in the present study was primarily driven by the clustered founder effect of F7, and remarkably, all the afore-mentioned CP24 deficiency phenotypes were manifested in further measurements on the F7 founder line, which had remarkably low CP24 expression levels, protein accumulation, super-complex abundance, and in vivo antenna size. Taken together, these findings suggest a strong contribution of CP24 expression level variation to the chromosome 10 QTL effect, although in the absence of further fine-mapping, it is not possible to rule out that any additional effects could also play a role.

### Utilization of identified QTL for improving photosynthetic efficiency in maize

Fv/Fm and ΦPSII are robust indicators of the maximum quantum yield and the operational efficiency of PSII photochemistry respectively (Murchie and Lawson, 2013; Baker, 2008). ΦPSII is tightly linked to the quantum yield of CO_2_ fixation (ΦCO2) measured by gas exchange in maize (Fryer et al., 1998; Pietrini and Massacci, 1998; Leipner et al., 1999) and other C4 species (Cousins et al., 2002; Ripley et al., 2007) across a range of environmental conditions. This suggests that the QTL for ΦPSII on chromosomes 9 and 10 are quite likely to impact the efficiency of photosynthetic CO_2_ assimilation. Although NPQ is less directly related to CO_2_ assimilation, it has been shown to transiently limit ΦCO_2_ in C3 species like *Nicotiana tabacum* (Kromdijk et al., 2016) and *Glycine max* (De Souza et al., 2022). Several recent studies have demonstrated variation in NPQ induction and relaxation in C3 species (Tom P.J.M. Theeuwen et al., 2022; Rungrat et al., 2019; Cowling et al., 2022)). Direct comparisons of NPQ kinetics between these studies and ours are complicated by the use of different phenotyping protocols and the use of different methods for modelling the response of NPQ to illumination and darkness. Despite this, it seems that NPQ is generally slower to relax in the aforementioned C_3_ studies compared to NPQ in the C_4_ crop maize in this study and (Sahay et al., 2023). Although substantial variation remains, this may suggest that NPQ kinetics are generally already closer to optimal in maize. Nevertheless, manipulation of photoprotection via enhanced NPQ amplitude may also have benefits for yield (Hubbart et al., 2018) and intrinsic water use efficiency (Głowacka et al., 2018) independent from NPQ kinetics.

For the identification of QTL or SNPs in collections of highly homozygous accessions in diversity panels or mapping populations to be relevant for improvement of photosynthesis in new maize cultivars, the associations need to remain significant in heterozygous backgrounds. Maize is a model species for heterosis (Shull, 1948). Indeed, plant breeding vastly enhanced maize yield by exploiting the positive effects of heterosis on plant size and growth rate. However, despite the importance of heterosis, its molecular basis and underlying genetic mechanisms are still largely unclear, despite some recent progress (Xiao et al., 2021). The level of heterosis varies strongly between traits and determines the ability to predict hybrid performance based on the inbred parent phenotypes, where traits with greater heterosis show relatively weaker correlations (Flint-Garcia et al., 2009). Evidence for heterosis effects on photosynthetic traits is inconsistent. Recent work in *A. thaliana* found heterosis effects on photosynthetic traits to be negligible across four sets of reciprocal F1 hybrids from contrasting ecotypes, despite strong heterosis effects on biomass accumulation (Liu et al., 2020). In maize, some work indicates that heterosis may impact photosynthetic traits to a significant degree (Mehta and Sarkar, 1992; Meena et al., 2021), whereas others report much less impact (Kamphorst et al., 2022). The creation of homozygous lines in naturally outcrossing species like maize gives rise to inbreeding depression, due to the accumulation of deleterious mutations. Indeed, maize inbred lines were shown to segregate for many SNPs predicted to result in deleterious variants (Mezmouk and Ross-Ibarra, 2014) for which complementation in the hybrid state could contribute to the observed hybrid vigor. While the importance of deleterious mutations for the QTL identified here is unclear, these may have played a role. Future research will therefore need to include validation of the QTL effects in a heterozygous state.

## Materials and Methods

All data analysis and plotting were conducted within R (R Core Team, 2021) if not stated otherwise. Raw sequencing reads are available on ENA under the study accession ID PRJEB67515. Phenotypic data presented here are available on electronic data archive library (Link provided upon publication of paper). Code associated with data analyses can be found on GitHub (https://github.com/capleo/MAGIC).

### Plant Material and Field Experiments

The MAGIC maize population described in (Dell’Acqua et al., 2015) was used in this study. This population was developed by crossing eight inbred lines (A632, B73, B96, F7, H99, HP301, Mo17 and W153R) and following a funnel breeding design. A ninth line (CML91) was introduced into the design as a two-way hybrid (B73xCML91) to account for failures in crossing B96xHP301. For this study, we used a subset of 320 RILs, different from those in the original paper, that well-represent the genetic composition of the population (Supplemental Figure S7).

Field experiments were carried out in 2021 and 2022 at the National Institute of Agricultural Botany on heavy clay loam soil types (NIAB, Cambridge, UK). The field experiment sites were approximately 500 m from each other across the two years (Supplemental Figure S1). Environmental data (precipitation, temperature, and light intensity) were collected from the same weather stations across the two years (Supplemental Figure S2).

In each year, 320 RILs were grown in an alpha-lattice experimental design with two replicates of 40 blocks each, where each block contained eight plots. The experimental design was constructed using the design.alpha function of the R/agricolae package (Mendiburu and Simon, 2015). Each plot consisted of four rows of ten plants representing a single RIL. The rows were spaced 25 cm apart and each plant within a row was 13 cm apart. In both years, seeds were sown by hand at a depth of 10-15cm. 301 RILs were common between each year (Supplemental Table 1.)

The 2021 experiment was sown on 13/05/2021 and the 2022 experiment was sown on 04/05/2022. Irrigation inputs were provided for the 2022 experiment only (Supplemental Figure S2). In both years, pesticides and fertilizers were applied throughout the experiment as necessary and according to manufacturer instructions (Supplemental Table S11).

### Chlorophyll fluorescence phenotyping

Six plants per RIL were measured, with three random plants coming from the representative plot in the first replicate of the alpha-lattice experimental design and three random plants coming from the second replicate. Measurements of chlorophyll fluorescence were performed on the leaf subtending the ear between three-to-eight days post silking. Measurements were spread out across three weeks in each year to account for the variation in silking time.

On each measurement day, the leaf subtending the ear from three plants from all plots selected to be measured that day was excised at the base. The excision point was then immediately submerged in water. This was performed at dawn (∼0530h) within a 30-minute window. The leaves were then returned to the laboratory and recut under water to maintain the water column and left in a 50 ml falcon tube under stable conditions for seven hours. After this time, a 2cm x 4cm strip of tissue was cut from the middle of the leaf and placed on top of damp filter paper that was on top of a non-reflective glass plate as per (Ferguson et al., 2020). 60-70 cut tissue samples were placed together on one sheet of paper in a grid-like fashion. A reference map of each grid of samples was made using QR codes attached to each leaf to cross reference the data with the sample it was generated from. Once all samples had been laid out, a non-reflective glass plate was placed on top of the samples to keep them flat and in place and to ensure that they did not desiccate. Samples were then dark adapted overnight using aluminum foil before performing chlorophyll fluorescence measurements in the following morning at 0900h. This experimental routine allowed us to perform additional measurements on the same leaf material prior to excising the tissue for chlorophyll fluorescence. We have recently demonstrated that the data generated for NPQ kinetics and maximum PSII operating efficiency through this approach is identical to that produced by directly measuring leaves that are still attached to the plant in maize (Ferguson et al., 2023b).

Chlorophyll fluorescence measurements were performed using a closed chlorophyll fluorescence imaging system (FluorCam FC 800-C, PSI, Czech Republic). Initially, the measuring beam was switched on to estimate minimal chlorophyll fluorescence (*Fo*). A saturating light pulse was then used to calculate dark-adapted maximum fluorescence (*Fm*). The actinic light source (1500 µmol m^-2^ s^-1^ photosynthetic active radiation (PAR)) was then switched on for 10 minutes. During these 10 minutes, a series of 12 saturating pulses were performed to estimate maximal fluorescence (*Fm’*) at the following intervals (in seconds): 20, 40, 60, 120, 180, 240, 300, 360, 420, 480, 540, 600. The light source was then subsequently switched off for 12 minutes and a further series of eight saturating pulses were performed to again estimate *Fm’* at the following intervals: 20, 40, 60, 120, 180, 360, 540.

Using the above-described data, the maximum quantum efficiency of PSII photochemistry was calculated as *Fv*/*Fm* where *Fv* is the variable fluorescence between *Fm* and *Fo*. At each point of measurement following the initial dark-adapted saturating pulse, NPQ was calculated as (*Fm*-*Fm’*)/*Fm’* and PSII operating efficiency (ΦPSII) was calculated as (*Fm’*/*F’*)/ *Fm’* (Figure 1; (Murchie and Lawson, 2013)). If the Fv/Fm value for any sample was below 0.70, it was removed from the analyses.

The slope from a linear model describing NPQ as a function of time was used to characterize the initial (0-80 seconds) response of NPQ to actinic light (NPQ induction slope) Figure 1B). Exponential models were used to characterize the induction of NPQ in response to the actinic light being switched on (Equation 1; Figure 1C)), the relaxation of NPQ following the actinic light being switched off (Equation 2; Figure 1D)), and the recovery of ΦPSII following the turning of the actinic light (Equation 3; Figure 1F)).

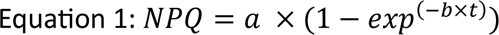

Where *a* represents the amplitude (NPQ induction amplitude), *b* represents the rate constant for the induction of NPQ (NPQ induction rate), and t represents time in seconds.

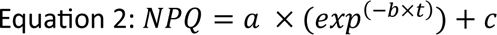

Where *a* represents the amplitude (NPQ relaxation amplitude) and *b* represents the rate constant for the induction of NPQ (NPQ relaxation amplitude). Here, *c* represents an offset to account for NPQ not reaching zero during this relaxation phase (NPQ relaxation offset).

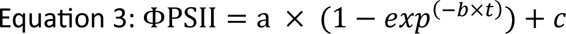

Where *a* represents the amplitude (ΦPSII recovery amplitude) and *b* represents the rate constant for the recovery of ΦPSII (ΦPSII recovery rate). Here, *c* represents an offset to account for a non-zero intercept (ΦPSII recovery offset).

We also extracted the maximum value for NPQ as well as the final values for NPQ and ΦPSII. Finally, we also calculated the photoprotection index (PI; Equation 4) which is a modification of the “pNPQ” approach of (Ruban and Murchie, 2012) and describes the proportion of the reduction in PSII efficiency is attributable to NPQ:

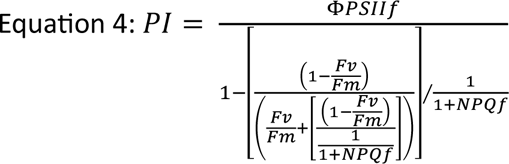

### Statistical Analyses

We extracted best linear unbiased predictors (BLUPs) for all measured traits from a general linear mixed model (Equation 5) performed via restricted maximum likelihood using the lmer and ranef functions from the lme4 R package (Bates et al., 2015) with the following formula with effects treated as random:

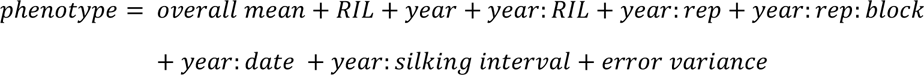

BLUP values were added to the population mean to generate predicted means for each trait. To extract year specific BLUPs, models identical to equation 5 were performed but without the year terms and including only data from either 2021 or 2022. Broad sense heritability (H2B) for each trait was calculated from the coefficients from the corresponding models as the ratio of genotypic variance to phenotypic variance according to the standard method (Schmidt et al., 2019).

Correlations between identical traits across 2021 and 2022 were tested for statistical significance via Pearson correlation analyses using the cor.test function in R. Correlations between all possible pairwise trait interactions on a year-by-year basis were also tested in this manner. One-way analysis of variance (ANOVA) comparison of means tests were performed to test for significant differences in the variance of each measured trait across 2021 and 2022 using the aov function in R.

### DNA extraction, genotyping and bioinformatic analysis

Genomic DNA of founders and RILs was extracted bulking young leaves of three seedling per genotype with the GenElute Plant Genomic DNA Miniprep Kit (Sigma Aldrich, Germany) following the manufacturer’s protocol. Integrity was assessed with agarose gel electrophoresis; DNA samples were quantified with the Qubit fluorometer (Invitrogen, Thermo Fisher Scientific, US). Samples were delivered to IGA Technology Services (Udine, Italy) to perform genotyping with the Single Primer Enrichment Technology (SPET), a protocol using single primer extension reactions to enrich pre-defined target loci. The set of SPET probes was developed starting from MAGIC maize founder haplotypes to maximize evenness of representation (Scaglione et al., 2019). Libraries were prepared with the Illumina TruSeq DNA Protocol (Illumina Inc., San Diego, CA, United States) and targeted fragments were sequenced with V4 chemistry in paired-end 150-bp mode on an Illumina HiSeq 2500 (Illumina Inc., San Diego, CA, United States). After initial quality assessment (FastQC), raw reads were filtered with erne-tool (ERNE2 package, version 2.1.1, http://erne.sourceforge.net/; (Fabbro et al., 2013). Trimmed reads were then mapped against the *Z. mays* reference genome (RefGen V4, Jiao et al., (2017)) with the BWA-MEM algorithm (Li and Durbin, 2009). The HaplotypeCaller algorithm (GATK), was used for variant call (Ryan Poplin et al., 2018). Genotyping was completed with GenotypeGVCFstool (Danecek et al., 2011) to derive raw single nucleotide polymorphisms (SNPs). SNPs were filtered using a Phred-scaled score greater than 30 and a SNP missingness per individual and locus lower than 20%.

### RNA sequencing data

Raw transcriptome data of eight founder lines except B96 were retrieved from ArrayExpress (https://www.ebi.ac.uk/arrayexpress/) under accession number E-MTAB-3173. Briefly, these data were generated on the MAGIC founder lines grown under controlled growth conditions: 24 °C, 55% relative humidity, 170μmol m^−2^ s^−1^ photosynthetically active radiation at plant level in a 16h/8h day/night cycle. Tissue for RNA extraction was sampled at the fourth leaf from three biological replicates, during the steady state growth phase. Samples were extracted with Trizol (Invitrogen), library prepared and sequenced with three technical replicates as detailed by (Baute et al., 2016). Raw reads were subjected to QC with erne-filter tool (ERNE2 package) and mapped against RefGen V4.0 (Jiao et al., 2017) using STAR v. 2.5.3a (Dobin et al., 2013) in 2-pass protocol, with default parameters (Miculan et al., 2021). Transcript abundances were extracted and normalized using *r/edgeR* (Robinson et al., 2010). For QTL where founder effects were distinct between B73 and F7, we additionally queried gene expression difference between these two founders utilizing the RNAseq datasets described in (Lee Cackett et al., 2023).

### Mapping of QTL and candidate gene identification

The genetic map was derived by anchoring the intermated B73 x Mo17 (IBM) genetic position over the SNPs derived after QC on the RefGen V4. Missing genetic distances were interpolated using values proportional to the physical distance. The reconstruction of the founder’ haplotypes in the RILs genomes (i.e., estimation of genotype probabilities) was executed using a Hidden Markov Model (HMM), implemented in R/qtl2 (Broman et al., 2019) that was used as platform for all the following mapping steps when not stated otherwise. QTL mapping of the phenotypes was performed using a mixed linear model incorporating kinship through the ‘Leave One Chromosome Out’ (LOCO) method. The genome scan regressions LOD scores were estimated using the sum of squared residuals for the null and alternative hypotheses. Thresholds of significance were defined based on 999 trait-specific permutations. The 95^th^ percentile of the permuted LOD distributions were set as threshold to identify peaks. For each of the significant LOD peaks, we used Bayesian credible interval estimation to define QTL confidence intervals, which were expanded to the closest SNP marker physical positions.

For each peak we estimated QTL coefficients within the confidence intervals, which represent parental haplotype effects as estimated by the model. The model conducts the mapping using parental haplotypes reconstructed on RIL genomes via HMM, and the haplotype effects estimates derive from phenotypic values observed in RILs carrying a given founder haplotype in the locus of interest. QTL coefficients in the confidence interval were clustered using a k-means algorithm with a function implemented in *r/fcp* (Hennig, 2010), that searched for the optimal number of clusters ranging from one to the number of founders except one. The most divergent groups of founder effects (clusters) were then used to test differential expression of the gene models within each corresponding confidence interval, using a generalized linear model on transcript count data derived from RNA sequencing. A threshold of FDR 0.1 was used for multiple testing correction. Due to the low number of RILs carrying the CML91 haplotype at the QTL, their effect as well as transcript abundances were excluded from these analyses (Dell’Acqua et al., 2015). The search for differentially expressed genes, based on founders’ haplotype effects, was only applied on significant peaks having confidence intervals with a span of less than 30 Mbp. The candidate genes identified were used to find the most similar Arabidopsis ortholog based on putative protein sequence identity. The procedure was carried out using the freely available database in PhytozomeV13 (Goodstein et al., 2012) as well as Plaza Monocot v. 4.5 (Van Bel et al., 2018).

To further the characterization of the identified QTL, we employed a *r/qtl2* built-in procedure designed to test marker associations within specific regions. This approach collapses genotype probabilities of the RILs to the SNPs of the founder genotypes at a single QTL region. For this analysis, we retrieved a comprehensive whole-Genome Sequencing (WGS) SNP database which was made available for the founders of our population (Dell’Acqua et al., 2015). From this database we derived 4,474,859 homozygous, polymorphic SNP. With this resource, we imputed founder SNPs onto the reconstructed RIL genomes. This way we were able to test associations in a given QTL, using a standard mixed linear model regression that incorporates kinship with the aforementioned LOCO method.

To fully exploit the genetic features of the MAGIC maize population we realigned existing WGS short-read sequencing data produced on the founders of the population (Dell’Acqua et al., 2015) against RefGen V4 using BWA-MEM. Before alignment all reads were quality-controlled by Fastqc v0.12.1 (Andrews, 2010) and filtered with bbduk from bbtools v38.87 (Bushnell, 2015) to gain reads with a quality of at least 25 and minimum length of 35bp. Bam files were filtered for a minimal mapping quality of 30 and sorted and indexed with samtools v1.17 (Danecek et al., 2021). Mapping statistics were called with qualimap v.2.2.2-dev (García-Alcalde et al., 2012). Copy number variations (CNVs) were determined with cn.mops v1.46.0 package (Klambauer et al., 2012) based on read counts in 100K bp bins for each chromosome. This approach provides insights into structural variation as inferred CNV within each QTL.

### Additional chlorophyll fluorescence phenotyping of B73 and F7 founders

Characterization of chlorophyll fluorescence parameters related to PSII antenna size and efficiency was performed using the LCF fluorometer of LI-6800 Portable Photosynthesis System (LI-COR Biosciences, USA). OJIP induction kinetics elicited by a 600 ms square pulse of 20,000 µmol m^-2^ s^-1^ irradiance was monitored using the continuous fluorescence signal mode in order to achieve the best signal-to-noise and temporal response (∼1.6 MHz bandwidth detection response with 4μs). The relative change of fluorescence yield was derived from the ratio of the continuous fluorescence signal divided by the continuous actinic irradiance signal monitored by the photodiode used for pulse optical control (Loriaux et al., 2013). The boosted flash intensity promoted fluorescence rise from *F*_o_ level to maximal *F*_j_ level and the corresponding reduction of the primary PSII quinone acceptor (Q_A_) within the first few turnovers of PSII. The half rise time *t*_1⁄2_ defined as the time to rising from *F*_o_ to one half of (*F*_j_-*F*_o_) was used as a proxy of the rate of photochemical reduction of Q_A_ and the reciprocal of *t*_1⁄2_ provided a relative estimate of the effective antenna size of PSII. The measurement was carried out on single leaf that was dark adapted for at least 30 min.

### Protein analyses

Thylakoid membrane protein complexes were isolated according to (Järvi et al., 2011). Briefly, 1 g of frozen leaf material (fourth fully expanded leaf) was ground in ice-cold grinding buffer (50 mM HEPES/KOH, pH 7.5, 330 mM sorbitol, 2 mM EDTA, 1 mM MgCl_2_, 5 mM sodium ascorbate, 0.05% BSA, 5 mM NaCl, and 10 mM NaF) and filtered through two layers of Miracloth (Sigma Aldrich). The flow-through was spun down for 5 min at 5000 × g at 4°C, and the resulting pellet was resuspended in 2 ml shock buffer (50 mM HEPES/KOH, pH 7.5, 5 mM sorbitol, 5 mM MgCl_2_, and 10 mM NaF) and then in 2 ml storage buffer (50 mM HEPES/KOH, pH 7.5, 100 mM sorbitol, 10 mM MgCl_2_, and 10 mM NaF). Finally, the pellet was carefully resuspended in 100 µl storage buffer. All the steps were performed under dim light and on ice. The chlorophyll concentration was determined from a 5 µl sample in 100% methanol according to (Porra et al., 1989). Aliquots of the thylakoid fractions were diluted with ACA buffer (50 mM BisTris/HCl (pH 7.0), 750 mM ε-aminocaproic acid, 1 mM EDTA, 0.25 mg ml^−1^Pefabloc, and 10 mM NaF) to 1 µg chl µl^-1^ and stored at -70°C.

For western blot analysis, 2 µg chl of thylakoid samples were denatured for 5 min at 70°C in 2x Laemmli sample buffer (138 mM Tris/HCl (pH 6.8), 6 M urea, 22.2% (v/v) glycerol, 4.3% (w/v) SDS, 5% (v/v) 2-mercaptoethanol, and 0.05% bromophenol blue). Samples were briefly spun down (5000 × g for 5 min), loaded onto a 12% Mini-PROTEAN® TGX™ Precast Protein Gel, and run in a Mini-PROTEAN® Electrophoresis Cell (Bio-Rad) using a Tris/glycine running buffer (25 mM Tris base, 190 mM glycine and 0.1% SDS). Proteins were then blotted onto a PVDF membrane (pore size 0.2 µm) using a Trans-Blot® Turbo™ Transfer System (Bio-Rad) and the “Mixed MW” protocol (7 min blotting time). The membrane was briefly washed in TBS buffer and then blocked with 10% non-fat milk in T-TBS buffer for one hour on a shaker. After that, the membrane was washed two times for 5 min with T-TBS on a shaker and incubated in the primary antibody (anti-CP24, Agrisera, 1:2500 in 1% milk in T-TBS) over night at 4°C, shaking. The next day, the membrane was washed four times for 5 min with T-TBS and incubated in the secondary antibody (Agrisera, goat anti-rabbit IgG, HRP conjugated, dilution = 1:12500 in 1% milk in T-TBS) for one hour on a shaker. Finally, the membrane was washed three times with T-TBS and twice with TBS for 5 min each and incubated with Clarity Western ECL substrate (Bio-Rad). After 5 min, the blot was imaged using a G:Box Chemi XRQ system (Syngene) with an exposure time of 3.5 min.

For blue native-polyacrylamide gel electrophoresis (BN-PAGE), 10 µg chl samples were solubilized with an equal volume of 2% digitonin (in ACA buffer, final concentration = 1%) by gently shaking samples at room temperature for 10 min. Afterwards, samples were spun down for 20 min at 18000 × g at 4°C and the supernatant was transferred into new tubes containing 1/10 of the volume of BN-PAGE sample buffer (100 mM BisTris/HCl (pH 7.0), 0.5 M ACA, 30% (w/v) sucrose, and 50 mg ml^−1^ Serva Blue G). After careful mixing, samples were loaded onto a 3-12% Native PAGE Bis/Tris precast gel (Invitrogen) and were run using the XCell SureLock™ Mini-Cell system (Invitrogen) on ice. The outer chamber was filled with clear anode buffer (50 mM BisTris/HCl, pH 7.0), whereas the inner chamber contained blue cathode buffer (50 mM Tricine, 15 mM BisTris/HCl (pH 7.0), and 0.01% Serva Blue G). Electrophoresis was performed using the following protocol: 75V for 30 min, 100 V for 5 min, 125 V for 30 min, 150 V for one hour and 175 V for 60-90 min. After the first 90 min, the blue cathode buffer was replaced with a clear cathode buffer, omitting Serva Blue G. The gel was then imaged using a flat-bed scanner.

### Analysis of CCD1 copy number by RT-qPCR

Real-time quantitative PCR (RT-qPCR) was used to determine the copy number of CCD1 in each founder. Genes of known single copy, VP14 (Tan et al., 2017) and ADH1 (Broothaerts et al., 2008) were used as references. Genomic DNA of founders was extracted from young leaves using the CTAB method. To improve the accuracy of the RT-qPCR, genomic DNA was first digested to completeness with EcoRI. Reactions were then prepared using 200 ng digested DNA, 200 nmole of forward and reverse primer sets for reference genes (ADH1,5’-CGTCGTTTCCCATCTCTTCCT-3’ and 5’-CCACTCCGAGACCCTCAGTC-3’; VP14, 5’-GCTGGCTTGGCTTGTATACTCTG-3’ and CCATCAGTCATATACTGTGAACAAATGT-3’) and CCD1 (5’-GGGAAGAGGGTGATGAAGTTGT-3’ and 5’-TGATATCCATTCACCTTGTCCAAA-3’) and 5 µl of SsoAdvanced Universal SYBR Green Supermix (172-5270; BioRad, Hercules, CA, USA). All samples were analyzed in triplicate using the CFX connect Real-Time PCR Detection System (1855201, BioRad, Singapore). The following RT-qPCR conditions were used: 3 min at 98°C, 40 times (10s at 98°C, 30s at 60°C), followed by a melting curve from 60°C to 95°C. Copy number was estimated using the ΔΔCt method (Livak and Schmittgen, 2001) and Ct values were normalized against B73 as a single copy reference (Tan et al., 2017).

## Acknowledgements

This work was supported by the European Union’s Horizon2020 research and innovation programme (No.862201) project CAPITALISE. We thank Haidee Philpott and Dr Tally Wright for consultation on experimental design and statistical analyses. We acknowledge the contribution of Vincent Forester, Tamanna Jithesh, Jordan Johnson, Simon Kapp, Ina Kruger, and Matthew Perry for their assistance in managing the field experiments and performing the phenotyping.

## Author Contributions

JNF and LC contributed equally to this manuscript. JK, MD, TL, JNF, and LC designed the experiments. JNF designed the field trials and carried out the phenotyping with assistance from JM. LC performed the genetic mapping analyses with assistance from SM and MST. All other data analyses were performed by JNF and LC with assistance from KS and RV. LM and BG performed the *in vivo* analyses of effective PSII antenna size. GT performed the *ccd1* copy number analyses. JW performed the protein analyses of CP24 in B73 and F7. Lee Cackett provided and queried the B73-F7 transcriptome dataset. JNF, LC, MD, and JK wrote the paper with input from all other authors.

## Supplemental Material

Supplemental Figure S1. Comparison of selected parameters across the two experimental years. (A-H) Boxplots demonstrating population wide variation across both experimental years for final ΦPSII, NPQ induction (slope from linear model), NPQ induction amplitude, NPQ relaxation amplitude, NPQ relaxation residual, Final NPQ, ΦPSII recovery amplitude, ΦPSII recovery residual. (I-P) Scatter plots demonstrating associations between the same traits across each experimental year. Significant differences and correlations are denoted at the following levels: * 0.05, ** 0.01, *** 0.001, **** 0.0001.

Supplemental Figure S2. Environmental parameters across the two experimental growing seasons. (A) Photosynthetically active radiation (PAR) in 2021. (B) Temperature in 2021. (C) Precipitation in 2021. (D) PAR in 2022. (E) Temperature in 2022. (F) Water inputs (precipitation and irrigation in 2022). Days where irrigation was applied are highlighted with asterisks.

Supplemental Figure S3. Broad-sense heritability for the main measured and modelled traits as derived from the 2021 (orange), 2022 (blue), and joint-year (mint green) linear mixed effects models.

Supplemental Figure S4. QTL mapping of ΦPSII at different timepoints within the induction (light) and relaxation (phases). Mapping is performed using the predicted means derived from the joint-year linear mixed effect models. (A) QTL for ΦPSII 20 seconds after the actinic light is switched on. (B) QTL for ΦPSII 60 seconds after the actinic light is switched on. (C) QTL for ΦPSII 20 seconds after the actinic light is switched off. (D) QTL for NPQ 560 seconds after the actinic light is switched off. (G) Broad-sense heritability (H^2^_B_) for ΦPSII over time.

Supplemental Figure S5. Overview of genes identified within all QTL intervals with previously demonstrated roles in the light dependent photosynthetic reactions, detoxification of reactive oxygen species (ROS), the Hatch-Slack Pathway (C4 photosynthesis), and the Calvin-Benson-Bassham Cycle (C3 photosynthesis). Genes are positioned approximately within the sites of these pathways for which they contribute to associated processes.

Supplemental Figure S6. Summary of results fast rise fluorescence kinetics of F7 and B73. (A) Differences in *F*_o_. (B) Difference in *F*_m_. (C) Differences in OJIP transients.

Supplemental Figure S7. Principal Components Analysis of the whole genotyped population (552) based on SPET genotyping SNPs (74,706): (A) PC1 vs PC2 (B) PC1 vs PC3. While founders are highlighted with different colors and larger dots, RILs include in this study are highlighted in light pink.

Supplemental Table S1. Predicted means for all main measured and modelled traits.

Supplemental Table S2. Predicted means for NPQ and ΦPSII at each measured timepoint.

Supplemental Table S3. Associations between all pair-wise trait interactions for predicted means derived from the 2021, 2022, and joint-year linear mixed effects model. Above-diagonals: Pearson correlations coefficients. Below diagonals: *P*-values. Significant (*p* < 0.05) interactions are highlighted in red.

Supplemental Table S4. Genes underlying all identified QTL

Supplemental Table S5. Genes where expression variation is associated with QTL haplotype effects

Supplemental Table S6. CCD1 copy number for the founder accessions of the MAGIC population.

Supplemental Table S7. Copy Number Variations (CNVs) identified within the QTL regions of the joint model.

Supplemental Table S8. BLAST results for Zm0001d048314

Supplemental Table S9. Genes underlying QTL in this present study and identified as candidate genes in Sahay *et al*. (2023)

Supplemental Table S10. Commonality between genes within the QTL intervals identified across the two experimental years in this present study and the genes identified as candidate genes across the experimental years in Sahay *et al*. (2023)

Supplemental Table S11. Agronomic inputs during the 2021 and 2022 field seasons

Supplemental Movie 1. QTL for NPQ during the light phase of the experimental protocol using predicted means derived from the joint-year mixed effects model.

Supplemental Movie 2. QTL for ΦPSII during the light phase of the experimental protocol using predicted means derived from the joint-year mixed effects model.

Supplemental Movie 3. QTL for NPQ during the dark phase of the experimental protocol using predicted means derived from the joint-year mixed effects model.

Supplemental Movie 4. QTL for ΦPSII during the dark phase of the experimental protocol using predicted means derived from the joint-year mixed effects model.

